# Positive allosteric modulation of metabotropic glutamate receptor 5 modulates Akt and GSK3β signaling *in vivo*

**DOI:** 10.1101/718700

**Authors:** Kari A. Johnson, P. Jeffrey Conn

## Abstract

**Background:** Positive allosteric modulators (PAMs) of metabotropic glutamate receptor 5 (mGlu_5_) have behavioral effects predictive of antipsychotic activity in experimental models such as amphetamine-induced hyperlocomotion (AHL). However, the signaling mechanisms that contribute to the antipsychotic-like properties of mGlu_5_ PAMs are not well understood.

**Methods:** Because the Akt/GSK3β pathway has been implicated in schizophrenia and is modulated by known antipsychotic drugs, we evaluated the effects of systemic administration of two mGlu_5_ PAMs on Akt and GSK3β signaling using western blot analysis in both naïve and amphetamine-treated adult male rats.

**Results:** In the dorsal striatum, the mGlu_5_-selective PAM VU0092273 (30 mg/kg) significantly increased Akt phosphorylation at residues associated with enhanced kinase activity, Thr308 and Ser473. Inhibitory phosphorylation of GSK3β at Ser9 was also increased. Similar effects were observed with a second mGlu_5_ PAM, VU0360172 (56.6 mg/kg). VU0092273 increased Akt phosphorylation levels in amphetamine-treated rats. Effects on Akt/GSK3β signaling were not limited to the striatum, as VU0092273 also increased Akt/GSK3β phosphorylation in the medial prefrontal cortex.

**Conclusions:** These findings suggest that mGlu_5_ PAMs that have antipsychotic-like efficacy in rats affect signaling pathways that are modulated by known antipsychotics, and raise the possibility that inhibition of GSK3β might contribute to the antipsychotic-like effects of mGlu_5_ PAMs.

## 1. Introduction

Dopamine and glutamate neurotransmitter systems are heavily implicated in the pathophysiology of schizophrenia. Positive symptoms of schizophrenia (i.e. psychosis) are associated with augmented dopamine synthesis and release, particularly in the associative striatum (reviewed in (McCutcheon et al., 2019). In addition, all currently approved typical and atypical antipsychotic drugs, which are primarily effective for treatment of positive symptoms of schizophrenia, have some affinity for dopamine receptors (Shin et al., 2011). However, these drugs are relatively ineffective at improving debilitating negative symptoms (e.g., flat effect, social impairment) or cognitive symptoms (e.g., deficits in executive function, working memory, and behavioral flexibility), suggesting that additional neurochemical systems are likely involved. Because blockade of NMDA receptors can recapitulate or augment a range of schizophrenia symptoms, dysfunction of glutamatergic transmission, particularly within circuits involving the prefrontal cortex, are also implicated (Howes et al., 2015; Meltzer, 2017; Moghaddam and Javitt, 2012). Recent efforts to identify novel treatments for schizophrenia include strategies to modulate glutamate transmission. The metabotropic glutamate (mGlu) receptor family of G protein-coupled receptors (GPCRs) are prominent modulators of glutamate transmission in the central nervous system (Niswender and Conn, 2010), and thus have been frequently investigated in the context of schizophrenia etiology and drug development (Stansley and Conn, 2018).

Among the eight subtypes of mGlu receptors, mGlu_5_ has received considerable preclinical attention as a therapeutic target for treating all symptom clusters associated with schizophrenia (reviewed in (Foster and Conn, 2017). Genetic deletion of mGlu_5_ reproduces some preclinical phenotypes of subcortical hyperdopaminergia and NMDA receptor hypofunction including hyperlocomotion, disrupted prepulse inhibition, and cognitive impairment (Brody et al., 2004; Burrows et al., 2015; Gray et al., 2009; Kinney et al., 2003; Lipina et al., 2007). Conversely, positive allosteric modulators (PAMs) of mGlu_5_ have behavioral effects that are predictive of antipsychotic activity in experimental models including reversal of hyperlocomotion induced by amphetamine (Gregory et al., 2013; Noetzel et al., 2012; Rodriguez et al., 2010; Rook et al., 2015a; Schlumberger et al., 2009) or NMDA receptor hypofunction (Gregory et al., 2013; Liu et al., 2008; Rook et al., 2015b; Spear et al., 2011), as well as reversal of prepulse inhibition following early life PCP exposure or apomorphine injection (Gregory et al., 2013; Rook et al., 2015b). In addition, mGlu_5_ PAMs enhance several types of learning in normal animals (Ayala et al., 2009; Gregory et al., 2013; Rook et al., 2015b; Spear et al., 2011) and alleviate cognitive deficits in both neurodevelopmental and NMDA receptor hypofunction models of schizophrenia (Clifton et al., 2013; Darrah et al., 2008; Gastambide et al., 2012; Gastambide et al., 2013; Gilmour et al., 2013; Stefani and Moghaddam, 2010; Uslaner et al., 2009). Based on these preclinical findings, mGlu_5_ PAMs have been advanced to preclinical and early clinical development for the treatment of schizophrenia (Rook et al., 2015b; Sturm et al., 2018).

Potentiation of NMDA receptor function has been posited as mechanism by which enhanced mGlu_5_ activity could exert antipsychotic-like and pro-cognitive effects; however, recent studies employing an mGlu_5_ PAM that does not affect NMDA currents in rats provided evidence for a dissociation between NMDA receptor interactions and many behavioral effects of mGlu_5_ PAMs (Rook et al., 2015b). Alternatively, the ability of mGlu_5_ PAMs to reverse behaviors induced by a hyperdopaminergic state in subcortical regions suggests that mGlu_5_ PAMs may have effects on cellular signaling that oppose the effects of dopamine receptor activation. However, the mGlu_5_-mediated signaling events that might intersect with pathways downstream of dopamine receptors *in vivo* are not well defined.

In recent years, the serine-threonine kinase Akt and downstream signaling molecules such as glycogen synthase kinase 3β (GSK3β) have received increased attention for their potential involvement in schizophrenia. Multiple genetic studies have identified Akt1 and Akt3 polymorphisms associated with schizophrenia, at least one of which is correlated with cognitive deficits (Blasi et al., 2011), and decreased levels of Akt and GSK3β have been found in postmortem brain samples from schizophrenic patients as well as in lymphocytes of schizophrenic patients (Beaulieu, 2012; Emamian, 2012; Emamian et al., 2004; Freyberg et al., 2010; Kozlovsky et al., 2002; Kozlovsky et al., 2004). In addition, transgenic mice with reduced Akt function display behavioral phenotypes reminiscent of those observed in schizophrenic subjects (Howell et al., 2017; Siuta et al., 2010), and Akt/GSK3 signaling abnormalities have been observed and correlated with cognitive deficits in several experimental systems used to study schizophrenia (Nadri et al., 2003; Takagi et al., 2015; Tamura et al., 2016; Willi et al., 2013). Biochemical studies have found that a number of currently prescribed antipsychotic drugs increase Akt phosphorylation in the striatum and prefrontal cortex in rodents (Roh et al., 2007; Sutton and Rushlow, 2011a), pointing to the Akt pathway as a potential target of antipsychotic action. The serine-threonine kinase GSK3β is constitutively active and is primarily regulated through inhibitory phosphorylation at Ser9 by Akt and other protein kinases (Kaidanovich-Beilin and Woodgett, 2011). Like Akt, GSK3β activity is altered by treatment with antipsychotic drugs (Alimohamad et al., 2005; Pan et al., 2015; Pan et al., 2016; Pan et al., 2018; Roh et al., 2007; Sutton et al., 2007; Sutton and Rushlow, 2011a, b). Conversely, amphetamine and apomorphine reduce Akt activity and GSK3β phosphorylation via activation of D_2_ dopamine receptors (Beaulieu et al., 2005; Beaulieu et al., 2004; Beaulieu et al., 2007; Shi and McGinty, 2007). Pharmacological or genetic inhibition of GSK3β markedly reduces amphetamine-induced hyperlocomotion (Beaulieu et al., 2004; Kalinichev and Dawson, 2011; Urs et al., 2012), whereas transgenic mice expressing constitutively active forms of GSK3 display enhanced responses to amphetamine and increased susceptibility to mood disturbances (Polter et al., 2010). Collectively, these studies point to Akt/GSK3 signaling as an important mediator of dopamine-related behaviors and a common target of clinically relevant antipsychotic drugs.

In the current study, we evaluated the ability of mGlu_5_ PAMs administered at doses that have antipsychotic-like and pro-cognitive effects in rats to modulate Akt and GSK3β signaling in the striatum and prefrontal cortex. We report that *in vivo* administration of mGlu_5_ PAMs increases Akt and GSK3β phosphorylation in the dorsal striatum of naïve male rats. Furthermore, pretreatment with an mGlu_5_ PAM increases Akt phosphorylation in amphetamine-treated rats. Finally, we provide evidence that mGlu_5_ PAM-induced Akt and GSK3β phosphorylation is not restricted to the striatum, as similar effects were observed in the medial prefrontal cortex. Taken together, these studies suggest that activation of Akt and downstream inhibition of GSK3β represent a cellular mechanism by which mGlu_5_ PAMs could counteract the behavioral effects of subcortical hyperdopaminergia and produce pro-cognitive effects in the medial prefrontal cortex.

## 2. Materials and Methods

### 2.1 Animals

Male Sprague-Dawley rats were purchased from Taconic (Indianapolis, IN) and allowed to acclimate to the housing facility within the Vanderbilt Rat Neurobehavioral Core for ∼1 week prior to experimentation. Rats weighed 250-300 g at the time of study. Animals were maintained in accordance with the guidelines of the American Association for the Accreditation of Laboratory Animal Care under a 12 hour light/dark cycle (lights on 06:00 to 18:00) with free access to food and water. All experiments were performed during the light cycle, were approved by Vanderbilt University’s Institutional Animal Care and Use Committee, and conformed to guidelines established by the National Research Council Guide for the Care and Use of Laboratory Animals. All efforts were made to minimize animal suffering and the number of animals used.

### 2.2 Drugs

VU0092273 and VU0360172 were synthesized in-house as previously described (Noetzel et al., 2012). VU0092273 (30 mg/kg) and VU0360172 (56.6. mg/kg) were dissolved in 10% Tween 80, sonicated for 30-60 minutes at 37°C, and injected intraperitoneally (i.p.) as a microsuspension in a volume of 3 ml/kg. Doses of VU0092273 and VU0360172 were chosen based on previous studies demonstrating that these doses effectively reverse amphetamine-induced hyperlocomotion (Noetzel et al., 2012; Rook et al., 2015a). Amphetamine hemisulfate (1 mg/kg, corrected for salt mass) was dissolved in saline and dosed subcutaneously (s.c.) in a volume of 1 ml/kg.

### 2.3 Sample preparation and western blotting

Following the indicated drug treatment times (15-105 minutes), rats were anesthetized under isoflurane anesthesia, decapitated, and brains were rapidly removed and placed in a chilled brain matrix. Coronal slices (1 mm thick) were prepared using razor blades. Slices containing medial prefrontal cortex (mPFC) and anterior dorsal striatum were frozen on a metal surface that was pre-chilled on dry ice. mPFC (prelimbic and infralimbic regions) was dissected by hand using a scalpel blade. Micropunches of dorsolateral striatum were obtained using a blunted 13-gauge needle. Following dissection, samples were placed into a microcentrifuge tube on dry ice and stored at −80°C prior to homogenization.

Samples were manually homogenized in 25-50 μL buffer containing (in mM): Tris HCl, 50, pH 7.4; NaCl, 50; EGTA, 10; EDTA, 5; NaF, 2; Na3VO4, 1; supplemented with 1X Complete Mini protease inhibitor cocktail (Roche) and phosphatase inhibitor cocktails 2 and 3 (Sigma-Aldrich). Homogenized samples were centrifuged at 16,100 x *g* in a table top microfuge for 10 minutes at 4°C. Supernatent fractions were removed, placed in a fresh tube, and protein assays were performed to determine protein concentration (DC™ Protein Assay, Bio-rad Laboratories, Inc.). 20-30 μg of each sample was mixed with 2X Laemmli buffer (Bio-rad Laboratories, Inc.), and heated at 65°C for 5 minutes. Samples were separated by SDS-PAGE on pre-cast gels (Bio-rad), transferred to nitrocellulose membrane, blocked with Odyssey blocking buffer (LI-COR Biosciences), and incubated with primary antibodies recognizing phosphorylated or total levels of proteins overnight at 4°C with gentle agitation. The following primary antibodies were used: phospho-Akt Ser473 (Cell Signaling #4058, 1:500), phospho-Akt Thr308 (Cell Signaling #2695, 1:500), total Akt (Cell Signaling #2920, 1:1000), phospho-GSK3β Ser9 (Cell Signaling #9322, 1:500), and total GSK3α/β (Santa Cruz Biotechnology #sc-7291, 1:250). Membranes were then incubated with IRDye-conjugated secondary antibodies (IRDye 680 for phosphoproteins, IRDye 800 for total proteins, Rockland Immunochemicals) for one hour at room temperature with gentle agitation. All antibodies produced bands at the expected molecular weights (60 kDA for Akt, 51 kDa for GSK3α, 46 kDa for GSK3β). For the antibody recognizing total GSK3α/β, only the 46 kDa band was measured for analysis.

### 2.4 Data analysis and statistics

Signals were detected using an Odyssey Quantitative Fluorescence Imaging System (LI-COR Biosciences). This method allowed simultaneous detection of phosphorylated and total protein levels. Band intensities were quantified using LI-COR Image Studio software. For each sample, the ratio of phosphorylated protein to total protein was obtained. All phosphorylation ratios were then normalized to the average phosphorylation ratio of samples from vehicle-treated animals. Statistical analysis depended on the experimental design. Unpaired t tests were used to compare vehicle-vs. drug-treated groups at individual time points. For analyses assessing phosphorylation levels under multiple conditions (Fig. 1d-g, Fig. 2), statistical comparisons were made using a two-way ANOVA followed by post hoc Bonferonni or Tukey multiple comparisons tests. For the sake of clarity, in graphs with multiple time points, only the values for PAM-treated animals are shown. GraphPad Prism 7.0 was used to create graphs and perform indicated statistical analyses.

**Figure 1.**
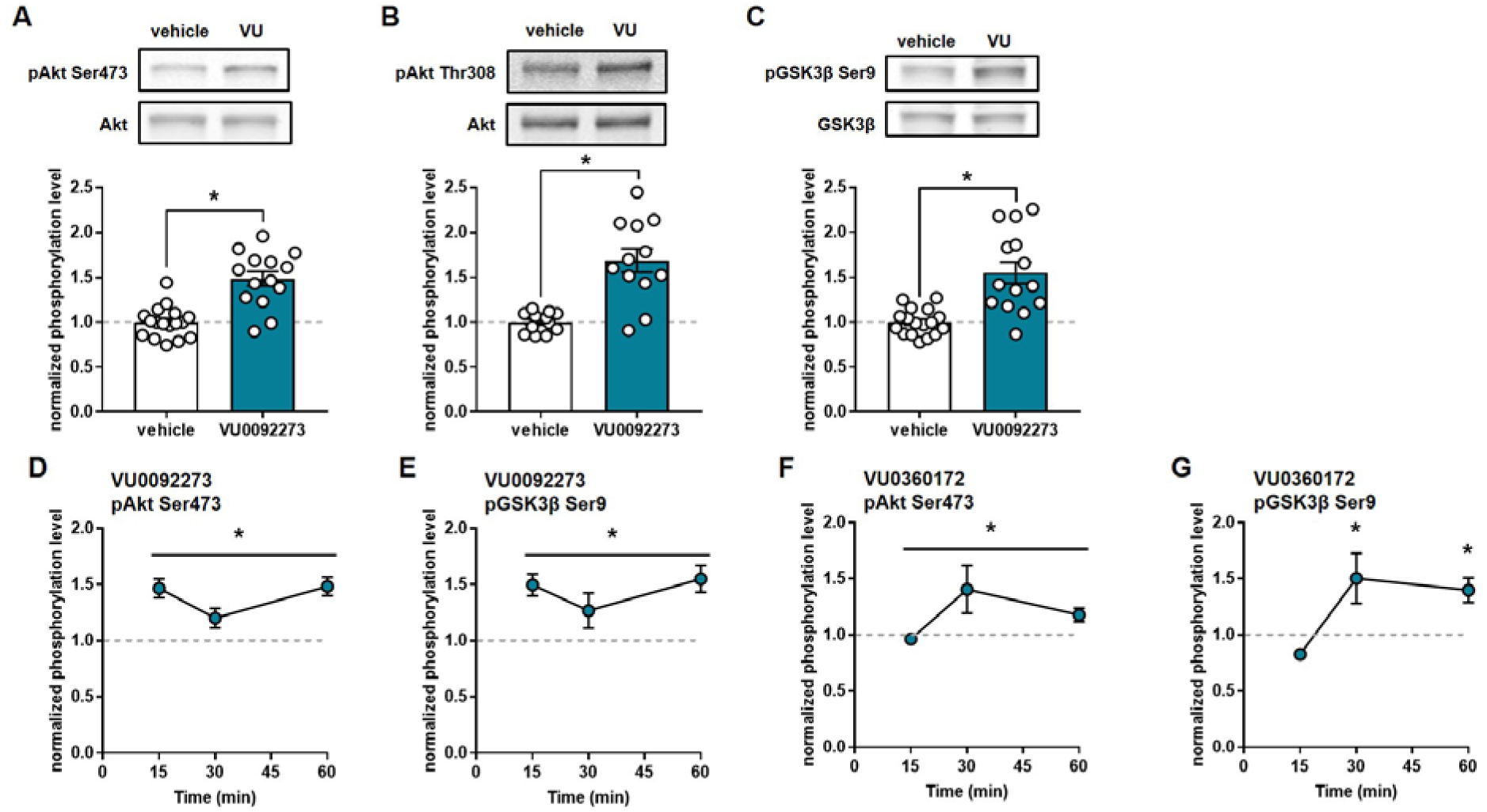
The mGlu5-selective positive allosteric modulators VU0092273 and VU0360172 increase Akt and GSK3β phosphorylation in the dorsal striatum. (a-c) Rats were given a single dose of VU0092273 (30 mg/kg, i.p., n = 17 [a and c] or 12 [b]) or vehicle (n = 14 [a and c] or 12 [b]) and were sacrificed one hour after injection. Phosphorylation levels of Akt Ser473 (a), Akt Thr308 (b), and GSK3β Ser9 (c) were measured in tissue punches from the dorsolateral striatum. Optical densities of phosphoprotein samples were first normalized to total Akt or GSK3β for each sample, then all samples were normalized to the mean value of the vehicle-treated group. Associated fluorescence images are representative examples of the phosphorylated (top) and total (bottom) bands for each protein. Asterisks (*) indicate significant differences between the vehicle and VU0092273 groups (p < 0.05, unpaired t test). (d-g) Drug-induced increases in pAkt Ser473 (d and f) and pGSK3β (e and g) when rats were euthanized at different times (15, 30, or 60 minutes) post-injection of VU0092273 (30 mg/kg, i.p., d and e, n = 7-14 rats/group) or VU0360172 (56.6 mg/kg, i.p., f and g, n = 7-14 rats/group). Optical densities for all samples were normalized to the mean value of vehicle treated controls (n = 5-17 rats/group). (d-f) Asterisks (*) indicate a main effect of PAM treatment analyzed by two-way ANOVA (p < 0.05). (g) Asterisks (*) indicate a significant difference between VU0360172 and vehicle at 30 and 60 minutes post-injection (p < 0.05, Bonferroni’s multiple comparisons test). All data represent the mean ± SEM.

**Figure 2.**
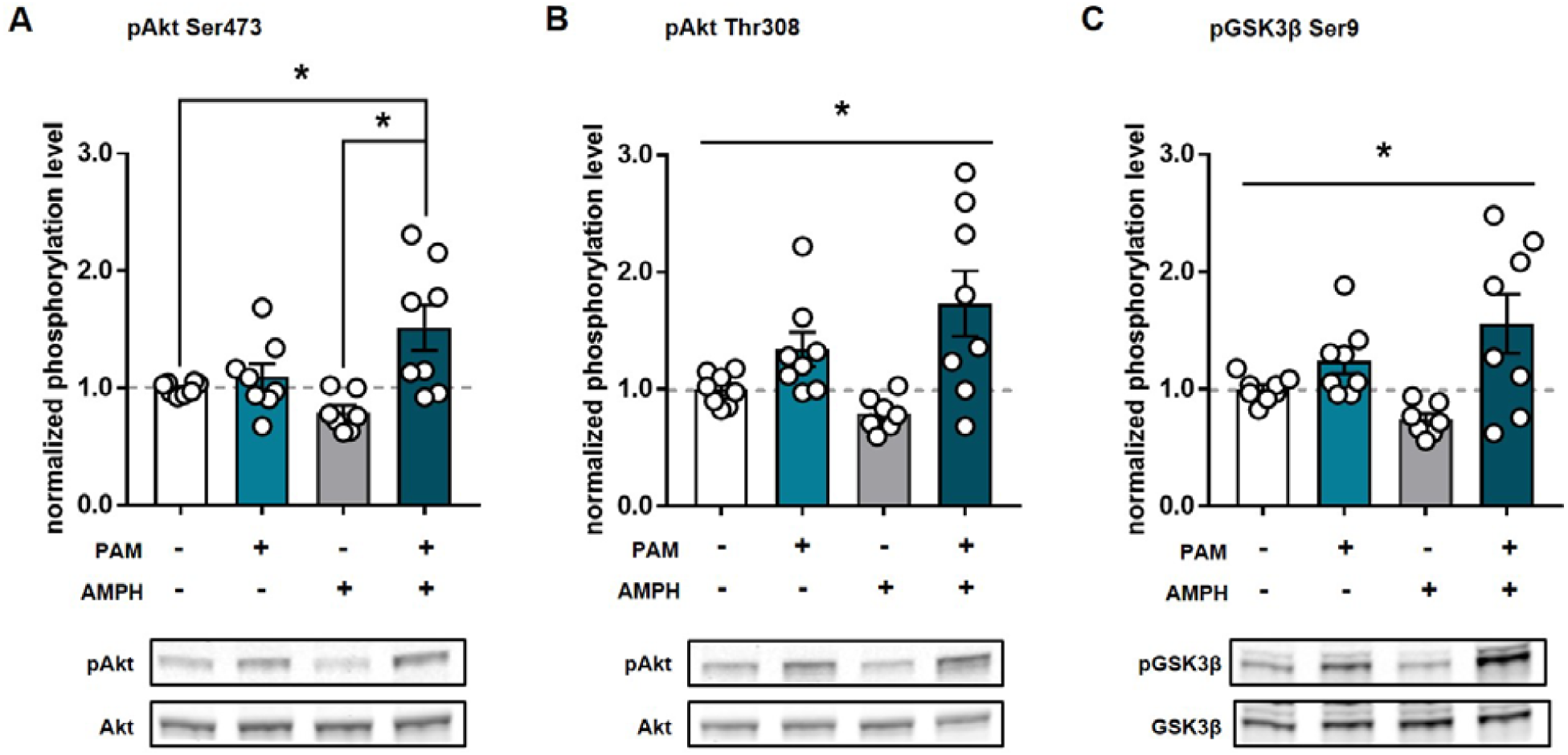
VU0092273 increases Akt phosphorylation in the dorsal striatum of amphetamine-treated rats. Rats were pre-treated with vehicle or VU0092273 (30 mg/kg), followed 15 minutes later by saline or amphetamine (1 mg/kg, s.c.; n = 8 for vehicle/saline, VU0092273/saline, and VU009273/amphetamine groups; n = 7 for vehicle/amphetamine group). Rats were sacrificed 90 minutes after amphetamine injection. Phosphorylation levels of Akt Ser473 (a), Akt Thr308 (b), and GSK3β Ser9 (c) were measured in tissue punches from the dorsolateral striatum. Optical densities of phosphoprotein samples were first normalized to total Akt or GSK3β for each sample, then all samples were normalized to the mean value of the vehicle-treated group. Associated fluorescence images are representative examples of the phosphorylated (top) and total (bottom) bands for each protein. (a) Asterisks (*) indicate a significant difference between the indicated groups (p < 0.05, Tukey’s multiple comparisons test). (b and c) Asterisks (*) indicate a main effect of VU0092273 treatment analyzed by two-way ANOVA (p < 0.05). All data represent the mean ± SEM.

## Results

### 3.1 The mGlu_5_ PAMs VU0092273 and VU0360172 increase Akt and GSK3β phosphorylation in the dorsolateral striatum

The ability of mGlu_5_ PAMs to reverse behavioral effects associated with a hyperdopaminergic state (e.g., amphetamine-induced hyperlocomotion) has been well established (Stansley and Conn, 2018). However, the mechanisms that underlie these effects are not well established. Because inhibition of Akt and subsequent activation of GSK3β have been implicated in the behavioral effects of amphetamine, and pharmacological or genetic inhibition of GSK3β attenuates amphetamine-induced hyperlocomotion (Beaulieu et al., 2004; Kalinichev and Dawson, 2011; Urs et al., 2012), we evaluated the ability of the mGlu_5_ PAM VU0092273 to modulate Akt and GSK3β signaling *in vivo*. We systemically administered VU0092273 and then measuring Akt and GSK3β phosphorylation in protein extracts from dorsolateral striatum samples. One hour after administration of the minimum dose that produces a maximal reversal of amphetamine-induced hyperlocomotion (30 mg/kg i.p.)(Rook et al., 2015a), VU0092273 significantly increased Akt phosphorylation at both Ser473 and Thr308 (pSer473: 1.49 ± 0.08 fold over vehicle, p < 0.0001; pThr308: 1.69 ± 0.13 fold over vehicle, p < 0.0001) (**Fig. 1a**,**b**). Concordantly, inhibitory phosphorylation of GSK3β phosphorylation at Ser9, a site that is known to be phosphorylated by Akt, was also increased (1.55 ± 0.12 fold over vehicle, p < 0.0001) (**Fig. 1c**). We then evaluated phosphorylation of Akt (at Ser473) and GSK3β in dorsolateral striatum samples at additional times points following administration of VU0092273 or a second mGlu_5_ PAM, VU0360172 (56.6 mg/kg, i.p.). For VU0092273 effects on pAkt, two-way ANOVA revealed a main effect of PAM treatment (F_(1,58)_ = 40.67, p < 0.0001) but not time (F_(2,58)_ = 1.91, p = 0.16), and there was no significant PAM x time interaction (F_(2,58)_ = 1.91, p = 0.16). For pGSK3β, we observed main effect of PAM treatment (F_(1,58)_ = 30.64, p < 0.0001), but not time (F_(2,58)_ = 1.04, p = 0.36) and no PAM x time interaction (F_(2,58)_ = 1.04, p = 0.36) (**Fig. 1d**,**e**). For VU0360172 effects on pAkt, two-way ANOVA revealed a main effect of PAM treatment (F_(1,54)_ = 5.64, p = 0.021) but not time (F_(2,54)_ = 2.41, p = 0.10), and there was no significant PAM x time interaction (F_(2,54)_ = 2.41, p = 0.10). For pGSK3β, we observed main effect of PAM treatment (F_(1,54)_ = 6.99, p = 0.011), a main effect of time point (F_(2,54)_ = 4.50, p = 0.016) and a significant PAM x time interaction (F_(2,54)_ = 4.50, p = 0.016). *Post hoc* comparisons of PAM vs. vehicle at each time point showed that pGSK3β was elevated at both 30 and 60 minutes post-injection (15 minutes, p > 0.99; 30 minutes, p = 0.0097; 60 minutes, p = 0.011 (**Fig. 1g**).

### 3.2 The mGlu_5_ PAM VU0092273 increases Akt phosphorylation in the dorsolateral striatum of amphetamine-treated rats

Treatment of rats or mice with amphetamine has been shown to decrease striatal Akt phosphorylation at Thr308 and thus increase GSK3β activity by reducing inhibitory phosphorylation at Ser9 (Beaulieu et al., 2005; Beaulieu et al., 2004; Beaulieu et al., 2007; Shi and McGinty, 2007). We therefore tested the ability of pretreatment with an mGlu_5_ PAM to reverse the previously reported biochemical effects of amphetamine on the Akt/GSK3β pathway. Rats were pretreated with vehicle (10% Tween 80) or VU0092273 (30 mg/kg, i.p.) 15 minutes prior to treatment with saline or amphetamine (1 mg/kg, s.c.), and samples were collected 90 minutes later and evaluated for pAkt (Ser473 and Thr308) and GSK3β (Ser9) phosphorylation. For all phosphorylation sites, two-way ANOVA revealed a main effect of PAM treatment (pAkt Ser473: F_(1,27)_ = 12.07, p = 0.0017); pAkt Thr308: F_(1,27)_ = 14.61, p = 0.0007; GSK3β: F_(1,27)_ = 13.36, p = 0.001) (**Fig. 2**). We did not observe a main effect of amphetamine treatment for any phosphorylation site (pAkt Ser473: F_(1,27)_ = 0.81, p = 0.38; pAkt Thr308: F_(1,27)_ = 0.31, p = 0.58; GSK3β: F_(1,27)_ = 0.0.3, p = 0.85). For pAkt Thr308 and pGSK3β Ser9, PAM x amphetamine interactions did not reach significance (pAkt Thr308: F_(1,27)_ = 3.20, p =0.085; GSK3β: F_(1,27)_ = 3.99, p = 0.058). For pAkt Ser473, there was a significant PAM x amphetamine interaction (F_(1,27)_ = 6.91, p = 0.01). *Post hoc* comparisons revealed significant elevations in pAkt Ser473 in rats treated with amphetamine and VU0092273 compared with rats that did not receive VU0092273 or amphetamine (p = 0.020). pAkt Ser473 was also elevated in rats treated with amphetamine and VU0092273 compared with rats treated with amphetamine alone (p = 0.001) (**Fig. 2a**). In this experiment, we did not observe significantly higher phosphorylation levels of pAkt Ser473 in rats treated with VU0092273 alone compared with rats that did not receive VU0092273 or amphetamine (p = 0.93). A likely explanation for the lack of effect of VU0092273 alone in this experiment is that more time had elapsed between drug injection and sample collection (105 minutes) compared with the time points at which we reported significantly higher levels of Akt and GSK3β phosphorylation (15-60 minutes) (**Fig. 1d**,**e**).

### 3.3 mGlu_5_ PAMs increase Akt and GSK3β phosphorylation in the medial prefrontal cortex

The results of our studies in the dorsal striatum implicate mGlu_5_ as a regulator of Akt and GSK3β signaling in a brain region that is associated with hyperdopaminergic states during psychosis (McCutcheon et al., 2019), and these neurochemical effects could contribute to mGlu_5_ PAM-mediated reversal of the behavioral effects of amphetamine. However, mGlu_5_ PAMs also enhance cognition in tasks related to prefrontal cortical networks, and downstream modulation of Akt/GSK3β signaling could contribute to pro-cognitive effects as well. Thus, we measured Akt and GSK3β phosphorylation in the mPFC (prelimbic and infralimbic regions) one hour after VU0092273 injection. Compared with effects observed in the dorsolateral striatum, VU0092273 caused a more modest elevation in pAkt Ser473 levels in the mPFC (1.18 ± 0.06 fold over vehicle, p = 0.018) (**Fig. 3a**). Increased pGSK3β levels were similar in magnitude to those observed in the dorsolateral striatum (1.50 ± 0.15 fold over vehicle, p = 0.003) (**Fig. 3b**).

**Figure 3.**
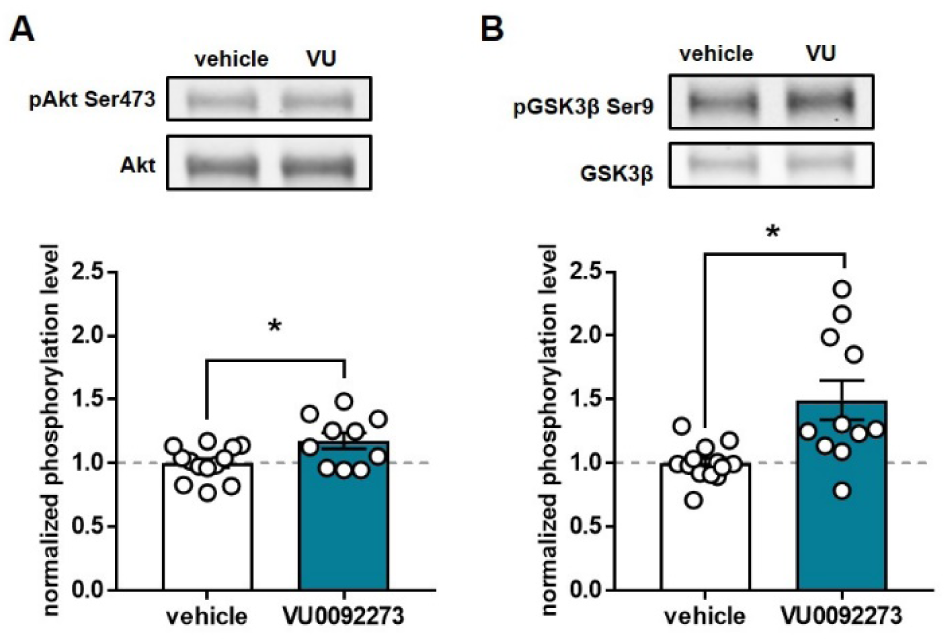
VU0092273 increases Akt and GSK3β phosphorylation in the medial prefrontal cortex. Rats were given a single dose of VU0092273 (30 mg/kg, i.p., n = 10-11) or vehicle (n = 13) and were sacrificed one hour after injection. Phosphorylation levels of Akt Ser473 (a) and GSK3β Ser9 (b) were measured in samples from the prelimbic and infralimbic regions of the medial prefrontal cortex. Optical densities of phosphoprotein samples were first normalized to total Akt or GSK3β for each sample, then all samples were normalized to the mean value of the vehicle-treated group. Associated fluorescence images are representative examples of the phosphorylated (top) and total (bottom) bands for each protein. Asterisks (*) indicate significant differences between the vehicle and VU0092273 groups (p < 0.05, unpaired t test). All data represent the mean ± SEM.

## 4. Discussion

The present study demonstrates that modulation of Akt and GSK3β signaling occurs in both the dorsolateral striatum and the mPFC following systemic administration of mGlu_5_ PAMs. Accordingly, mGlu_5_ PAMs join a growing list of drugs with known or predicted antipsychotic effects that inhibit GSK3β signaling in the striatum, likely through an Akt-dependent mechanism (reviewed in (Beaulieu, 2012; Freyberg et al., 2010). These include typical antipsychotics such as haloperidol, which block D2 receptors in the striatal complex (Sutton et al., 2007); aripiprazole, a D2 receptor partial agonist (Pan et al., 2015; Pan et al., 2016); atypical antipsychotics such as olanzapine, quetiapine, and clozapine, which display varying degrees of D2 antagonism and 5-HT2A receptor antagonism (Sutton and Rushlow, 2011a; Xi et al., 2011); and group II mGlu receptor agonists, which have preclinical behavioral profiles predictive of antipsychotic effects (Sutton and Rushlow, 2011b). In addition, various drugs that have clinical utility as antidepressants (e.g. fluoxetine, ketamine) and mood stabilizers (e.g. lithium, valproate) also lead to inhibition of GSK3β (Beaulieu, 2012; Dandekar et al., 2018), indicating that GSK3β inhibition is a common feature of a remarkable variety of psychotropic drugs that have demonstrated therapeutic efficacy in a broad range of psychiatric disorders. Our finding that VU0092273 increases Akt phosphorylation in amphetamine-treated rats supports the idea that mGlu_5_ PAMs could also counteract hyperdopaminergic striatal states through this pathway.

Although GSK3β inhibition has not been confirmed as a critical signaling event underlying the therapeutic effects of antipsychotic drugs, studies performed in rodents in which GSK3 isoforms are genetically altered to increase or reduce their activity provide several lines of evidence that GSK3β inhibition may play an important role in the mechanisms of action of these drugs. For example, mice with heterozygous deletion of the gene encoding GSK3β display a marked reduction in amphetamine-induced hyperlocomotion (Beaulieu et al., 2004), and this is mimicked by GSK3β deletion specifically in D2-expressing neurons (Urs et al., 2012). Concordantly, GSK3 inhibitors reduce amphetamine-induced hyperlocomotion (Beaulieu et al., 2004; Kalinichev and Dawson, 2011). Conversely, transgenic mice that express constitutively active forms of GSK3α and GSK3β exhibit enhanced hyperlocomotion in response to amphetamine, suggesting that GSK3 activity plays a permissive role in the behavioral expression of amphetamine-induced hyperlocomotion (Polter et al., 2010). Together, these studies identify inhibition of GSK3, and specifically GSK3β, as a downstream mechanism by which mGlu_5_ activation could reverse behaviors elicited by hyperdopaminergic activity in the striatum.

Our findings indicate that in addition to modulation of striatal Akt signaling, mGlu_5_ PAMs also increase Akt and GSK3β phosphorylation in the medial prefrontal cortex, an area associated with cognitive deficits in schizophrenia. Interestingly, functional MRI studies have indicated prefrontal cortical hypoactivation associated with a human genetic variant of GSK3β that is correlated with schizophrenia diagnosis (Blasi et al., 2013). In rodents, GSK3β has been shown to regulate AMPA and NMDA receptor function in PFC neurons (Khlghatyan et al., 2018; Wang et al., 2013; Xi et al., 2011). In addition, GSK3 inhibitors increase PFC theta oscillations and PFC-hippocampus coherence at doses that have pro-cognitive effects (Nguyen et al., 2017). Thus, mGlu_5_ modulation of the Akt/GSK3β pathway has the potential to impact PFC physiology to produce pro-cognitive effects. Further studies will be needed to directly link mGlu_5_ PAM actions on physiology and cognition with signaling via Akt or GSK3β. In addition to actions in PFC circuits, mGlu_5_ modulation of hippocampal function, or PFC-hippocampus interactions, might also contribute to mGlu_5_ PAM-mediated enhancement of cognition in normal animals and in experimental systems used to study schizophrenia. Akt signaling is required for forms of mGlu_5_-mediated synaptic plasticity in the hippocampus that are frequently associated with impaired cognition (Hou and Klann, 2004). A recent study reported that after repeated dosing, the mGlu_5_ PAM VU0409551 increases Akt and GSK3β phosphorylation in the hippocampus of wild-type mice and reverses lower phosphorylation levels in the hippocampus of serine racemate knockout mice, which have impaired NMDA receptor function and a schizophrenia-like phenotype (Balu et al., 2016). Rescue of Akt/GSK3β signaling in the hippocampus is associated with reversal of deficits in synaptic plasticity and contextual fear memory, suggesting that mGlu_5_ effects on Akt/GSK3β signaling in the hippocampus could be an additional substrate for pro-cognitive effects of mGlu_5_ PAMs.

In conclusion, we have found that mGlu_5_ PAMs modulate Akt and GSK3β signaling *in vivo*, which could contribute to the antipsychotic-like behavioral effects of these drugs, particularly in terms of their ability to reverse amphetamine-induced hyperlocomotion. Further studies will be necessary to determine a causal role of Akt/GSK3β pathway modulation in the anti-hyperdopaminergic and pro-cognitive effects of mGlu_5_ PAMs. Our increased understanding of the influence of mGlu_5_ PAMs on signaling pathways *in vivo* could guide future drug development towards compounds that are optimized to impact critical downstream effector proteins and therefore provide maximal therapeutic benefit.

## Acknowledgements

This work was supported by National Institutes of Health grants R01MH062646, R37-NS031373 (to P.J.C.) and F31 NS067737 (to K.A.J.).

## Contributions

P.J.C. and K.A.J. conceived the project and wrote the manuscript. K.A.J. performed experiments and analyzed data. The authors thank Dr. Jerri Rook for advice on experimental design.

